# Viro3D: a comprehensive database of virus protein structure predictions

**DOI:** 10.1101/2024.12.19.629443

**Authors:** Ulad Litvin, Spyros Lytras, Alexander Jack, David L Robertson, Joe Grove, Joseph Hughes

## Abstract

Viruses are intracellular parasites of organisms from all domains of life. They infect and cause disease in humans, animals and plants but also play crucial roles in the ecology of microbial communities. Tolerance to genetic change, high-mutation rates, adaptations to hosts and immune escape has driven high divergence of viral genes, hampering their functional annotation and phylogenetic inference. The protein structure is more conserved than sequence and can be used for searches of distant homologs and evolutionary analysis of divergent proteins. Structures of viral proteins are traditionally underrepresented in public databases, but recent advances in protein structure prediction allows us to address this issue. Combining two state-of-the-art approaches, AlphaFold2-ColabFold and ESMFold, we predicted models for 85,000 proteins from 4,400 human and animal viruses, expanding the structural coverage for viral proteins by 30 times compared to experimental structures. We also performed structural and network analyses of the models to demonstrate their utility for functional annotation and inference of distant phylogenetic relationships. Taking this approach, we examined the deep evolutionary history of viral class-I fusion glycoproteins, gaining insights on the origins of coronavirus spike protein. To enable further discoveries, we have created *Viro3D* (https://viro3d.cvr.gla.ac.uk/), a virus species-centred protein structure database. It allows users to search, browse and download protein models from a virus of interest and explore similar structures present in other virus species. This resource will facilitate fundamental molecular virology, investigation of virus evolution, and may enable structure-informed design of therapies and vaccines.

## Introduction

Viruses are obligate intracellular parasites whose replication depends on the metabolism and translational machinery of cellular organisms. Viruses have the capacity to evolve rapidly and infect organisms from all domains of life. They play crucial roles in ocean biogeochemical cycles ^1,2^ and control of prokaryotic populations in the human gut microbiome ^3^ but also infect and cause disease in crops, livestock and humans. Virus particles are the most abundant biological entities on our planet ^4^ with metagenomic and metatranscriptomic studies only starting to sample the staggering genetic diversity of viral communities ^5–7^ .

Intracellular parasites, like viruses and mobile genetic elements, seem to be an inherent property of any replicating system ^8^. Viruses likely emerged on multiple independent occasions with the origin of the most ancient lineages probably coinciding with the origin of life and preceding the appearance of the last common ancestor of cellular organisms (LUCA) ^9^. The genome of LUCA likely already carried an early antiviral defence system ^10^ implying that cellular organisms are inseparable from viruses and have been locked in a continuous arms race for the last 4 billion years. Independent evolutionary origins of viruses are reflected in the modern virus taxonomy by a specific taxonomic rank, realms, which brings together viruses that share a set of conserved genes usually involved in genome replication or virion formation ^11,12^.

The diversity of virus genome architectures with their frequent modular organisation ^13^, high mutation rates ^14^, positive selection, and dependency on host-cells drive viruses to evolve faster than cellular organisms. Viruses exchange genetic material with their hosts and other viruses, turning their genomes into a mosaic form where each section has a different evolutionary history ^15^. Countless examples of gene exchange between cellular organisms and viruses ^16^ and co-opting of proteins that evolve to fulfil a new function (called exaptations) ^17,18^ highlight the evolutionary importance of genetic exchange between viruses and their hosts.

However, frequent genome reorganisations and high levels of divergence make the identification of gene function, investigation of deep phylogenetic relationships and taxonomic assignments at higher ranks particularly challenging. In cases like these, comparison of protein structures for inference of evolutionary relatedness tends to be more reliable ^19^. Protein function is defined by tertiary structure which is, as a result, more conserved than nucleotide or amino acid sequence ^20^. Despite the striking diversity of viral sequences, viral protein structures are underrepresented in public databases. Experimental protein structures from a viral source constitute less than 10% of the Protein Data Bank (PDB) ^21^.

Recent advances in machine learning have made it possible to predict protein structures from sequence alone, achieving accuracy similar to that of experimental structure determination ^22–24^. These state-of-the-art approaches have been applied at scale to produce comprehensive databases of predicted protein structures. The AlphaFold Structural Database (AFDB) ^25^ contains more than 214 million models for proteins from UniProtKB ^26^ predicted using AlphaFold2 ^23^. However, viral proteins were excluded from the prediction efforts. Evolutionary Scale Modelling (ESM) Metagenomic

Atlas became the largest structural database with more than 600 million models for proteins from metagenomic samples (MGnify database ^27^) predicted using ESMFold ^24^. Although this database contains some viral structures, they come predominantly from viruses of prokaryotes and unicellular eukaryotes. The database also does not provide an intuitive way of searching for structures from a specific taxonomic group.

Systematic exclusion of viral proteins from structure prediction efforts and their underrepresentation in public databases created a gap that is currently being addressed by the scientific community ^28,29^. In our study, we have generated 170,000 viral protein structure predictions from 4,400 animal viruses using ColabFold and ESMFold. We assessed the model quality produced by both methods and performed structural analysis of the proteins to expand their functional annotation. We also demonstrate that protein structure can guide the inference of deep phylogenetic relationships between viruses, using class-I membrane fusion glycoproteins as an example. To meet the requirements of the virology community we created Viro3D, a fully-searchable and browsable database, allowing users to visualise and download models for all proteins from a virus of interest and explore similar structures present in other virus species (https://viro3d.cvr.gla.ac.uk/). We expect that this resource will find broad utility, accelerating fundamental molecular virology, enabling studies of virus evolution, and facilitating structure-informed development of therapies and vaccines.

## Results

### Systematic viral protein structure prediction with ColabFold and ESMFold

Our initial focus has been on predicting protein structures for human and animal viruses (Figure 1a). We relied on data from the International Committee on Taxonomy of Viruses (ICTV) Virus Metadata Resource (VMR), which provides a comprehensive list of virus species and representative isolates, along with their GenBank accession numbers and host associations. At the time of our analysis, the latest version of the ICTV VMR included 3,173 virus species infecting vertebrate and/or invertebrate hosts, represented by 4,407 virus isolates/genotypes. These viruses encoded a total of 71,274 proteins, spanning over 29.2 million amino acid residues (Figure 1b). To simplify the analysis and protein structure prediction procedure, we excluded polyproteins from the dataset replacing them with annotated mature peptides and protein regions that increased the total number of analysed protein records to 85,162.

**Figure 1.**
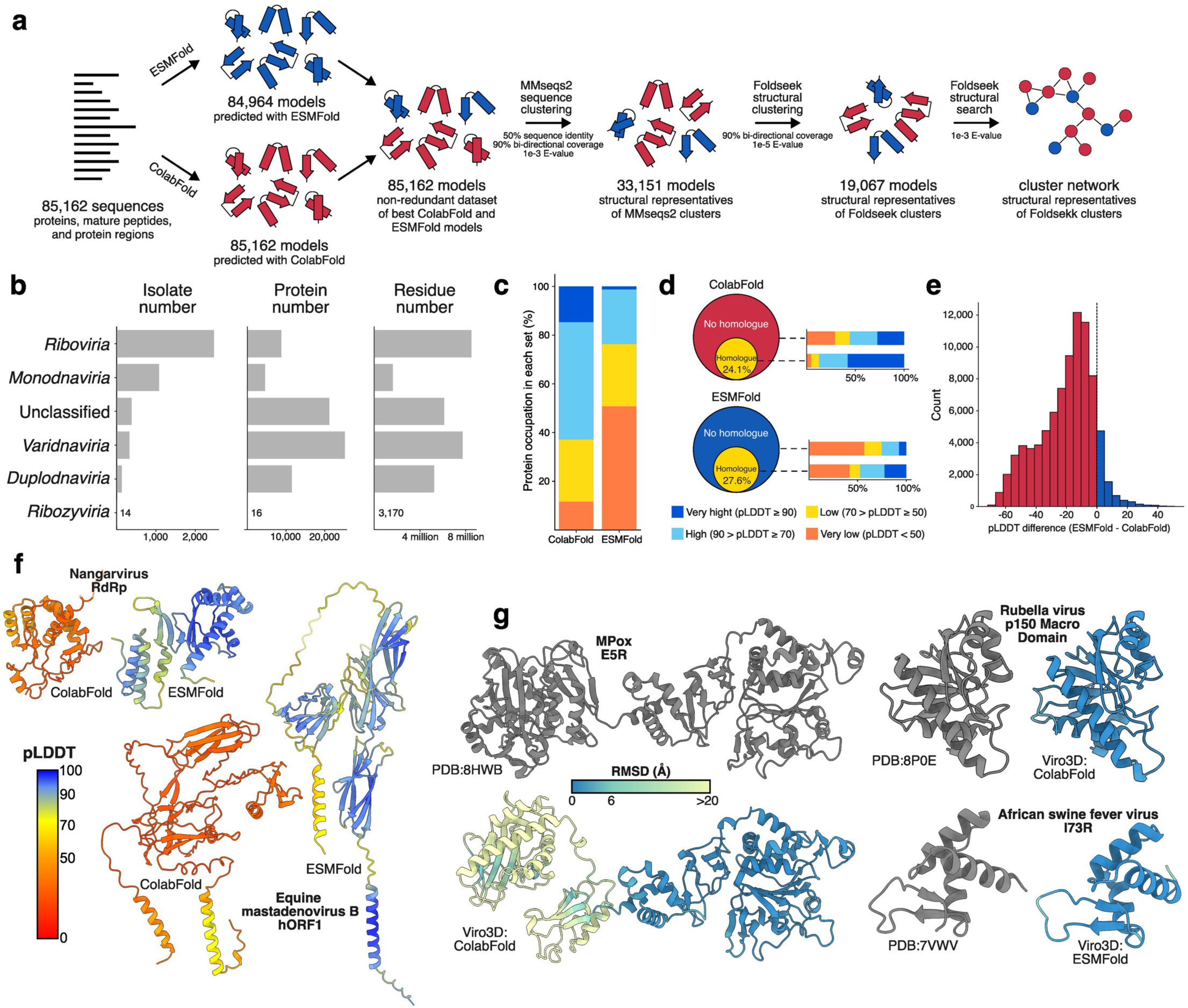
Expanding structural coverage of the human and animal virosphere. **a.** Summary of protein structure prediction, and downstream analyses (relevant to Figure 2). We predicted structures for 85,162 proteins, peptides and regions encoded by 4,407 viruses from the ICTV VMR, with vertebrate and/or invertebrate hosts, using ColabFold and ESMFold. A non-redundant set of models was composed using models with the highest average pLDDT from either ColabFold or ESMFold. These were subjected to sequence and structure-based clustering, as described in the text. **b.** Description of the target dataset with the number of virus isolates, proteins and residues per viral realm. Unclassified viruses were those lacking realm annotation (e.g. *Baculoviridae*) **c.** Percentage of ColabFold and ESMFold models with very high (dark blue), high (light blue), low (yellow) and very low (orange) average pLDDT score. **d.** Difference in the confidence of predictions by ColabFold and ESMFold that are covered by PDB sequence homologues (yellow area) and those lacking PDB homologues (red and blue area, respectively). Plots display proportion of total amino acid residues. **e.** Distribution of differences in average pLDDT scores between ColabFold and ESMFold models for each protein record where both models are available. Positive pLDDT difference values indicate that ESMFold model is better (blue bars), negative value – ColabFold model is better (red bars). **f**. Ribbon diagrams, color-coded by pLDDT, illustrating targets for which ESMFold produces a much more confident model. **g.** Recent experimental structures (grey), determined after the AlphaFold2/ESMFold training date cutoff, compared to their cognate Viro3D predicted structures, which are color-coded by root-mean squared deviation (Å) after rigid-body alignment with their experimental counterparts. Low RSMD values (blue) indicates excellent agreement.

For protein structure prediction, we applied two state-of-the-art approaches: ColabFold ^30^, a method based on AlphaFold2 ^23^ and dependent on multiple sequence alignments (MSAs), and ESMFold ^24^, a method that uses ESM-2, a transformer protein language model, and infers structure from input protein sequence alone. With ColabFold we successfully predicted structures for all 85,162 records (27.2 million residues, covering 93.1% of total amino acid residues). Due to compute limitations in predicting longer proteins, ESMFold yielded slightly fewer successful predictions – 84,964 protein records (27.0 million residues, covering 92.3% of amino acid residues). Structural coverage varied across viral realms, with most of them achieving coverage between 95% and 100% by both methods (Figure S1a). However, because of the lack of mature peptide and region annotation for large sections of *Riboviria* polyproteins, the coverage of this realm did not exceed 79.5% of total amino acid residues when we used ColabFold and 78.7% when we applied ESMFold. Nonetheless, since experimental structures in the PDB cover less than 3.3% of amino acid residues present in proteins of human and animal viruses (approximately 890,000 amino acids), we managed to expand the structural coverage for viral proteins by more than 30 times.

Overall, ColabFold models showed higher accuracy than those produced by ESMFold (Figure 1c, S1b). A total of 17.2 million residues predicted with ColabFold (63.3% of the modelled residues) were assigned high or very high quality, based on predicted local distance difference test (pLDDT) score. This number increased to 87.4% (5.7 million residues) when evaluating models where a sequence homologue was available in the PDB at the time of AlphaFold2 training, dropping to 55.7% (11.5 million residues) for regions without homologues (Figure 1d). In contrast, only 31.6% of residues predicted by ESMFold (8.5 million residues) achieved high or very high quality. Nonetheless, ESMFold models followed the same trend: with higher quality predictions for structures where a PDB sequence homologue was available at the time of training (47.6%, or 3.5 million residues) and lower quality for those without a homologue (25.5%, or 5 million residues). This suggests that training data may account for some higher accuracy models but, nonetheless, high confidence predictions can still be achieved for sequences that were not well represented on the PDB (this may be particularly important for viral proteins, which are under-represented in experimental structural data).

For 9.1% of the records (7,769 proteins), the ESMFold models demonstrated higher accuracy than the ColabFold models. Although these improvements were typically minor (Figure 1e), for 2.0% of the records (1,753 proteins), the difference in pLDDT scores exceeded 10 points (Figure 1f). One contributing factor to lower performance of ColabFold was low MSA depth (Figure S1c), confirming the importance of having sufficient similar sequences available in public databases. The protein records where ESMFold outperformed ColabFold also tended to have longer sequences (Figure S1d).

We sought to test the accuracy of predicted models through comparison to experimental structures solved after the AlphaFold2 and ESMFold training dates, and for which there were no sequence homologues available on the PDB at the time of training. Very few targets met these criteria; nonetheless, we found excellent agreement between predicted and experimental structures (Figure 1g), with the only major source of divergence being the positioning of, otherwise, well-folded domains (Figure S1e).

For all subsequent analysis, we created a non-redundant set of 85,162 models (Figure 1a) representing each protein record with either ColabFold or ESMFold model depending on which one has the highest confidence (average pLDDT score).

In parallel with our work, two other efforts have recently performed systematic structure prediction for viral proteins. Nomburg et al. predicted structures for 67.7K proteins from 4.4K eukaryotic viruses derived from RefSeq entries in the NCBI Viruses portal ^29^. The Big Fantastic Virus Database (BFVD) did not consider viral taxonomy but used viral sequences from UniRef30 clusters to generate 351.2K protein structures ^28^. We compared our Viro3D dataset to these using Foldseek (Figure S2a). There was moderate overlap with Nomburg et al.’s dataset with 47.3K structures being unique to Viro3D; whilst only 12.9K BFVD structures were shared with Viro3D (72.3K being unique to Viro3D). To understand this low overlap better, we performed proteome level comparisons for example human pathogens. Here, the majority of entries had a match in the Nomburg et al. dataset, but almost no proteins had a complete counterpart in BFVD, with many appearing only as truncated protein fragments (Figure S2b). Indeed, comparison of protein lengths for all entries in BFVD and Viro3D, demonstrated that short sequences predominate in BFVD (Figure S2c). This likely reflects the sequence selection and structure prediction strategy underlying BFVD ^28^. The high frequency of matches between Viro3D and the Nomburg et al. dataset permitted side-by-side comparison of model quality, assessed by pLDDT prediction confidence. Here, Viro3D models outperformed their counterparts in Nomburg et al. (Figure S2d & e). This is explained by the MSA generation method (drawing only on RefSeq virus sequences) and structure prediction workflow of Nomburg et al. ^29^. The benefits of Viro3D are well illustrated by comparisons of proteome structures from influenza A virus, where Viro3D provides complete high-confidence models for all proteins (Figure S2f). These results demonstrate that Viro3D is the most comprehensive database of high-quality protein structure predictions for human and animal viruses.

### Exploration of the viral protein structure space by clustering and network analysis

We started with clustering the non-redundant set based on 90% bi-directional sequence overlap and 50% sequence identity that resulted in 33,151 sequence clusters. Then we selected a representative for each cluster, taking the highest confidence model (by average pLDDT score), and clustered this set of structural representatives based on 90% bi-directional structure overlap and 1e^-5^ E-value of the structural alignment. This produced 19,067 structural clusters, 64.35% of which contained only one member (singleton clusters). Finally, we performed a Foldseek search among the representatives of these clusters to generate a structure similarity network of the viral proteins.

By applying restrictive clustering parameters, we ensured high structural homogeneity and consistency of functional annotation within each cluster (Figure S3) but allowed homologous viral proteins to form multiple structural clusters. For instance, one of the most abundant proteins in our dataset, RNA-dependent RNA polymerase (RdRp), formed at least 110 structural clusters. The structure similarity network allowed us to address this issue by capturing communities of clusters that possess the same general protein fold or share a protein domain.

To demonstrate the power of structural network analysis, we examined the distribution of various common and hallmark viral proteins. This includes proteins involved in genome replication: RdRp, Reverse Transcriptase (RT), and Family B DNA polymerase (PolB); virion morphogenesis: Single Jelly Roll (SJR), Double Jelly Roll (DJR), and HK97 major capsid proteins (MCP); and membrane fusion: class I, class II, and class III fusion glycoproteins (FG). Querying against individual experimental structures, representing each of these hallmark proteins, we performed Foldseek analyses across our structural network. This identified communities that match each of our hallmark protein groups (Figure 2a). Protein communities located in the centre of the network (e.g. RdRp, RT and SJR MCP) tend to be strongly interconnected, likely because of the clusters of polyproteins that possess multiple functional regions and therefore bring different communities together. This network-focussed structural search allowed us to find many more proteins than a similar search against the simple non-redundant set of 85,162 structures (Figure 2b), for example we achieved a 90% increase in the number of identified RdRp structures. Therefore, by leveraging the fundamentally conserved nature of structure over sequence, combined with clustering and network analysis, we can achieve extremely sensitive detection of deep evolutionary relatedness. This permits rapid and efficient navigation of viral protein structure space and allows mapping of protein form and function across diverse species.

**Figure 2.**
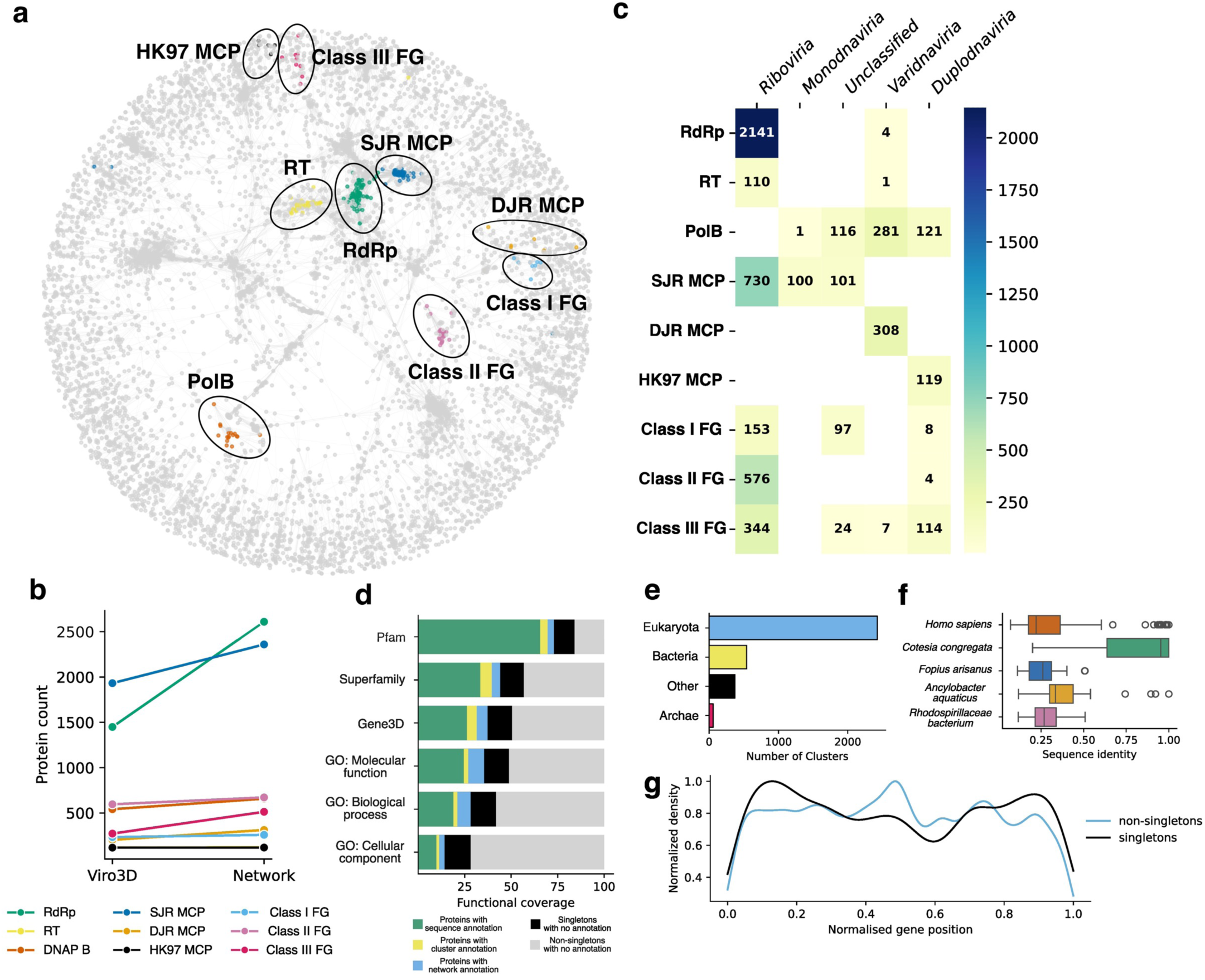
Cluster and network analysis of viral protein structures. **a.** Structure similarity network of viral proteins. Each node represents a cluster. Edges connect clusters that share structural similarity. Communities of hallmark proteins are highlighted in different colours. RNA-dependent RNA polymerase (RdRp), Reverse Transcriptase (RT), Family B DNA polymerase (PolB), Single Jelly Roll (SJR), Double Jelly Roll (DJR) and HK97 major capsid proteins (MCP), class I, class II, and class III fusion glycoproteins (FG). **b.** Number of viral protein structures found for each hallmark protein using a structural search against the Viro3D non-redundant dataset of 85,162 structures or against the network of structural clusters. **c.** Distribution of hallmark viral proteins across viral realms. Each square shows the number of viruses that possess structural homologues of hallmark proteins. **d.** Expansion of function annotation using structural information. Each bar shows a percentage of protein records per viral realm that possess Pfam annotation based on InterProScan (green), structural cluster expansion (yellow), structural network expansion (light blue). Percentage of protein records that do not have a Pfam annotation are black (if they belong to singleton clusters) or light grey (if they belong to non-singleton cluster). **e.** Number of viral protein clusters that have structural similarity to entries in the AFDB. Each bar represents the source of the AFDB structure. Structures coming from metagenomic, environmental and unclassified samples were labelled as other. **f.** Distribution of sequence identities between viral proteins and five cellular organisms from the AFDB with the highest number of structural matches. Box represents the interquartile range, the line inside the box represents the median value. Whiskers extend from the box to the smallest and largest values within 1.5 times the interquartile range. Outliers and are plotted as individual points. **g.** Distribution of genome positions of viral protein-coding genes. Proteins that belong to singleton clusters are in black, protein that belong to non-singleton clusters are in light blue.

We plotted the distribution of hallmark proteins across viral families (Figure S4) and realms (Figure 2c) successfully identifying RdRp or RT in 91% of *Riboviria* species. We also showed that five members of the *Varidnaviria* realm (*Poxviridae* and *Iridoviridae* families) possess an RT in addition to PolB. Four of these RTs are in a community of RdRp clusters (reflecting their shared ancestry) and therefore can be found using an RdRp probe, while one is clustered with other RTs. PolB is abundant in the realms *Varidnaviria*, *Duplodnaviria* and several viral families that currently do not belong to any realm (e.g. *Baculoviridae*). We also found a PolB in the *Bindnaviridae* family which belongs to the *Monodnaviria* realm. As for major capsid proteins, Double Jelly Roll MCP dominates in *Varidnaviria*, HK97 MCP in *Duplodnaviria*, while Single Jelly Roll MCP is prevalent in realms *Riboviria*, *Monodnaviria* and family *Baculoviridae*. All three classes of fusion glycoprotein are abundant in *Riboviria*, while class III is the dominant glycoprotein in *Duplodnaviria*. Class I and III FGs are also common among members of the *Baculoviridae* family.

We also used this approach to expand functional annotation, propagating sequence-based annotation using structural clusters and structural network. Out of 85,162 protein records, 65.6% have at least partial Pfam annotation (Figure 2d). By propagating Pfam annotations to unannotated cluster members, we expanded the functional coverage by 3.99% (3,395 records). The propagation of Pfam annotation to clusters that do not have any annotated members using the structural network expanded the functional coverage by an additional 3.37% (2,870 records, Figure S5). The consistency of propagated annotation is 100% for the majority (70.3%) of protein records (4,403 out of 6,265 records, Figure S6). We also expanded Gene3D and Superfamily annotation which gave an increase of 11.1% and 10.7% respectively. Expansion of gene ontology (GO) annotation for molecular function, biological process and cellular component increased the number of annotated records by 10.9%, 9.2% and 4.4% respectively. Therefore, clustering of structures and network analysis permits the identification and/or functional annotation of as-yet unclassified viral proteins. Nonetheless, it is important to note that despite propagation, there is still a high proportion of proteins with no known function (Figure 2d).

To estimate the number of protein structures shared between viruses and cellular organisms, we performed a structure similarity search between cluster representatives and the AlphaFold Structural Database (AFDB, Figure 2e). Consistent with the notion that viruses are a source of novel protein folds ^29^, the majority of viral proteins do not share detectable homology with cellular life, with only 17.8% of clusters (3,393 out of 19,067) having significant structural similarity to proteins in the AFDB. Of these homologues, 71.5% are coming from Eukaryota, 15.9% from Bacteria, 1.7% from Archaea and 10.9% from metagenomic and environmental samples. However, this number is likely to be an overestimation because in many instances hits are coming from endogenous or symbiotic viruses. For instance, many *Homo sapiens* hits with high sequence identity are proteins from the integrated *Human betaherpesvirus 6A* (Figure 2f), while hits from *Cotesia congregata* and *Fopius artisans,* two species of parasitoid/parasitic wasps, are proteins from symbiotic viruses of genera *Bracoviriform* and *Alphanudivirus*, respectively.

14.4% of the protein records form singleton clusters and potentially represent structural novelty. Our species-focussed approach captured the genomic context of all predicted structures and, therefore, allowed us to investigate the genome positions of singleton and non-singleton clusters (Figure 2g). Interestingly, unique protein structures are more frequently found at the start or end of viral genomes, while common protein structures (which are likely to be associated with critical functions) tend to have a more uniform distribution across the genome length. The picture is different for some viral realms (Figure S7), nonetheless, this suggests that genomic termini are hotspots for evolutionary innovation, which may drive the emergence of novel protein functions and adaptations to hosts.

### The deep evolutionary history of class-I fusion glycoproteins

Class-I fusion glycoproteins can be found in a wide range of important human pathogens including SARS-CoV-2, HIV, Influenza and Ebola, and have been exapted by mammals to mediate placental morphogenesis ^31^. Mechanistic knowledge of these proteins informs rational vaccine design ^32^. It is thought that class-I fusion glycoproteins share a common origin, however, extensive sequence divergence has all but erased this deep ancestry at the amino acid level. Therefore, classification of fusion glycoproteins is typically achieved through structural and functional characterisation. Indeed, recent experimental structures of retroviral Env class-I glycoproteins suggest shared ancestry with fusion glycoproteins of negative-sense RNA viruses ^33,34^; this evolutionary relationship is also supported by analysis of the viral ‘fossil-record’ provided by endogenous viral elements ^35^. Nonetheless, traditional structural biology and/or sequence analyses provides only a fragmentary picture of glycoprotein ancestry. We reasoned that ultra-sensitive homology detection, achieved through structural similarity and network analysis (Figure 2a & c), will permit a virosphere-wide survey of class-I fusion glycoproteins and provide a clear view of their deep-evolutionary history.

First, to gain a species-level perspective, we generated a structure-informed map of the human and animal virosphere. This allows simultaneous visualisation of all viruses represented in Viro3D, on to which the distribution of proteins can be projected. This was achieved by systematic structural comparison of each virus’ proteome (see Methods) resulting in a scatter plot, with viruses segregated by structural similarity (Figure 3a), which broadly recapitulates viral taxonomy (Figure S8).

**Figure 3.**
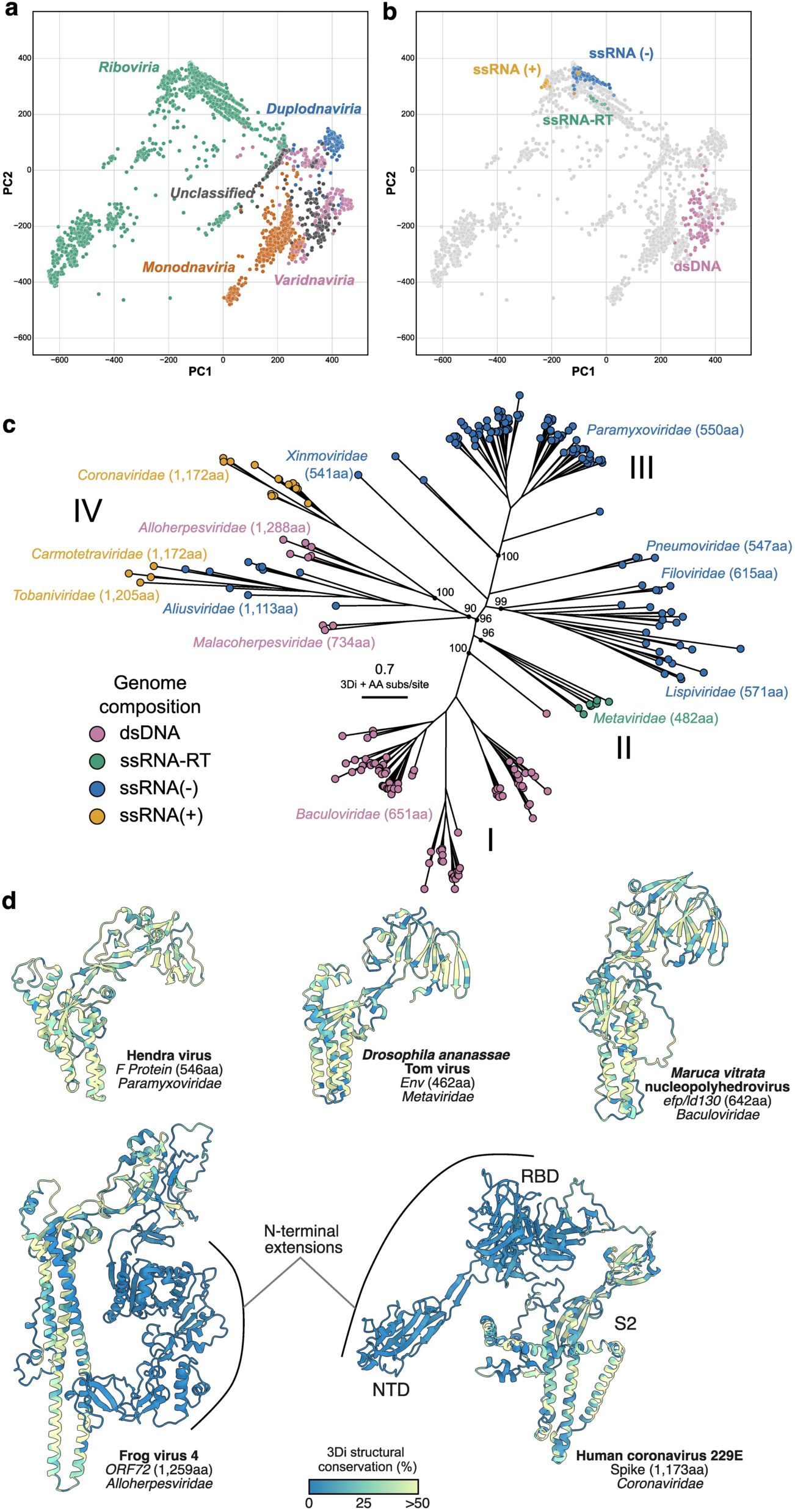
The deep evolutionary history of class-I fusion glycoproteins. **a.** Structure-informed map of the human and animal virosphere, each data point represents a virus in the Viro3D database (see Methods). Viruses are color-coded by realm. **b.** Distribution of class-I fusion proteins across the human and animal virosphere. Coloured data points represent viruses in which we detect a class-I fusion protein using structural searches and network analysis, with RSV-F as a reference. Viruses are colour-coded by genome composition, as labelled. **c.** Combined 3Di and amino acid phylogenetic inference of identified class-I fusion proteins, achieved by structure-guided sequence alignment using Famsa3di. Tips and labels are coloured by genome composition. Labels indicate viral genera and, in parentheses, the corresponding average length of glycoprotein. Major clades are labelled, as described in the main text. Node support values are provided at important branch points. **d.** Example structures; ribbon diagrams are colour-coded by structural conservation, assessed by the 3Di alignment, as denoted in the key. Labels indicate N-terminal extensions, present in clade IV glycoproteins, and pertinent regions of spike (N-terminal domain: NTD, receptor binding domain: RBD, and the S2 subunit). Note, predicted structures are monomeric, as opposed to the native trimeric state of class-I fusion glycoproteins, and domain positioning (e.g. NTD and RBD) may be inaccurate.

Mapping the viruses that possess class-I fusion glycoproteins (identified by structural homology to RSV-F, Fig. 2c) demonstrates a distribution across viral realms, indicative of extensive genetic exchange (Figure 3b). Alongside expected instances of class-I fusion glycoproteins in negative sense RNA viruses, reverse-transcribing viruses and positive sense Nidoviruses, we discovered previously unknown class-I fusion glycoproteins in the *Herpesvirales* (specifically, aquatic herpesviruses) and confirmed their presence in the *Baculoviridae*. Note that detection was achieved using a single structural reference (RSV F), we expect further instances may be found (for example in the *Retroviridae*) using alternative reference structures.

We performed structure-guided sequence alignment, permitting phylogenetic inference using both structure and sequence information, revealing a coherent evolutionary history (Figure 3c). This reveals four major clades (designated I-IV in Figure 3c). The first contains the *Baculoviridae*, where the class-I fusion system may, in fact, be more prevalent than the prototypical gp64 class-III fusogen previously characterised in baculoviruses ^36^. The second clade contains reverse transcribing viruses, from the *Metaviridae*, which share insect hosts with the *Baculoviridae*, providing a feasible route of genetic exchange ^37^. The third clade is monophyletic for the *Mononegavirales*, this may suggest a single genetic acquisition by an ancestral negative sense RNA virus, from which class-I fusion glycoproteins propagated throughout the order. The final clade (IV) contains a mixture of viral taxonomies including negative sense *Aliusvirdidae*, positive sense *Coronaviridae* and *Carmotetraviridae*, and dsDNA *Herpesvirales*. The clade IV glycoproteins are particularly long when compared to the rest of the phylogeny (most being >1100 residues; Figure 3c), suggesting adaptive diversification.

Comparison of structures from across the phylogeny reveals a central structurally-conserved architecture common to all of the identified class-I fusion proteins (Figure 3d) consistent with recent experimental studies ^33,34^. Example clade IV glycoproteins from an aquatic herpesvirus and a human coronavirus, however, possess very long N-terminal extensions; this includes the N-terminal domain (NTD) and receptor-binding domain (RBD) that constitute the majority of the S1 subunit in coronavirus spike. Moreover, the region conserved across all species broadly corresponds to the fusogenic S2 subunit of spike. Notably, the herpesviruses are basal in clade IV and the *Coronaviridae* form a high confidence branch with the *Alloherpesviridae* (Figure 3c). This suggests that the coronavirus spike glycoprotein originated from ancient genetic exchange with a dsDNA herpesvirus.

## Discussion

In this work, we expanded the structural coverage for viral proteins by 30 times compared to that of experimental structures by generating structural models for 85,000 proteins from 4,400 human and animal viruses. Approximately 64% of the produced protein models are confident predictions with an average pLDDT score above 70. Moreover, 65% of residues in the predicted dataset have high or very high confidence. We confirmed that, in general, ColabFold has a higher chance of producing a confident structure than ESMFold. However, since ESMFold models have higher confidence for almost 10% of protein records, do not require an MSA and are usually much faster to generate, the combination of both approaches is highly beneficial. We also demonstrated that Viro3D exceeds the coverage and quality of existing repositories of viral protein structure predictions.

By performing clustering of protein structures encoded by human and animal viruses the diversity of viral proteins was reduced to approximately 19,000 distinct protein structures; almost 65% of these proteins are unique within our dataset. Interestingly, in viruses with linear genomes (realms *Varidnaviria*, *Duplodnaviria* and, to some extent*, Riboviria*), these unique proteins are usually found closer to the ends of the genomes, which probably represent hot spots of gene acquisition or *de novo* gene origination.

More than 82% of distinct viral structures are unique to viruses and do not have apparent structural homologs in cellular organisms. This high percentage of unique viral structures may seem surprising given the extensive horizontal gene transfer between viruses and their hosts, and may suggest that even acquired proteins undergo extensive remodelling and diversification in the context of a viral genome. Therefore, we acknowledge that the number of viral proteins retaining more distant structural similarity to cellular components may be higher than that demonstrated in our analysis; indeed parallel studies have identified slightly higher numbers of cellular homologues in viruses ^28,29^.

We showed that structure similarity searches combined with a structural network allowed us to substantially accelerate the search process by increasing the number of homologous structures identified in a single run. For instance, using single RdRp and RT structures we identified RNA-directed polymerases for 90% of viruses in the *Riboviria* realm. This demonstrates that structural searches and network analysis are an extremely efficient means of traversing large evolutionary distances, even where underlying sequence homology may be negligible.

Since each viral realm represents an independent origin of an evolutionary distinct virus group ^38^, protein structures shared across multiple realms can be considered instances of horizontal gene transfer. Genome replication and capsid morphogenesis modules, commonly used for the taxonomic assignment of viruses at higher taxonomic ranks, tend to be hallmark features of specific realms and are rarely shared beyond. RdRp and RT are (almost) unique to members of *Riboviria*, while DJR MCP and HK97 MCP are specific to *Varidnaviria* and *Duplodnaviria*, respectively ^39^. PolB is distributed more broadly: realms *Varidnaviria*, *Duplodnaviria* and some unclassified DNA viruses. Most families in the *Monodnaviria* rely on rolling-circle replication endonuclease ^40^. We found PolB only in one family of this realm: *Bidnaviridae* ^41^. SJR MCP is also widely distributed, including RNA viruses within the realm *Riboviria* and viruses with circular dsDNA genome (realm *Monodnaviria* and unclassified family *Baculoviridae*) ^39^. Membrane fusion glycoproteins are common but not considered hallmark proteins, and appear to be more mobile being spread across multiple realms. Class III fusion glycoproteins seem to be the most widely distributed among human and animal viruses, being absent only in the realms *Monodnaviria* and *Ribozyviria*.

Fusion glycoproteins are fundamentally important for virus transmission, pathogenesis and spillover, and are major targets for host immunity. Viro3D has provided a new opportunity to comprehensively survey the distribution of glycoproteins and, using structure-guided approaches, infer their evolutionary past. Using just a single structural reference to query Viro3D (RSV F glycoprotein), we identified 259 highly-diverse class-I fusion glycoproteins, many sharing <10% sequence identity. This includes both known instances (e.g. in *Mononegavirales*) and newly discovered glycoproteins (e.g. in aquatic herpesviruses). Combined structure and sequence phylogenetics provided the first view of the deep evolutionary history of class-I fusion glycoproteins. This is consistent with multiple independent horizontal gene transfers from an, as yet unidentified, ancestral source.

Whilst all of the identified glycoproteins share a structurally conserved central fold, one clade in particular shows signs of extensive adaptive diversification. The phylogenetic topology of this clade suggests that coronaviruses gained their spike glycoprotein by genetic exchange with aquatic herpesviruses. These glycoproteins have doubled in size by gaining novel regulatory elements, such as the NTD and RBD in the S1 subunit of spike. This provides a unique evolutionary explanation for the membrane fusion mechanism of coronaviruses: the S1 subunit is proteolytically separated from spike and is shed following receptor engagement, leaving the structurally-conserved S2 to mediate fusion ^42^. Thus, to achieve virus entry coronaviruses undergo a regulated unmasking of an ancestral fusogen, which is fundamentally conserved across all class-I fusion glycoproteins. In summary, using class-I fusion glycoproteins as an exemplar, we have demonstrated that structure-guided discovery, enabled by Viro3D, is likely to provide unprecedent new insights on the origins and evolution of viruses.

Whether for lab-based investigators wishing to bring three-dimensional context to their molecular virology experiments, researchers performing structure-guided design of therapies, or computational biologists studying deep virus evolution, we expect the rich structural dataset presented here to be valuable resource for the virology community. Viro3D is fully searchable and browsable here: https://viro3d.cvr.gla.ac.uk/.

## Methods

### Dataset preparation

6,721 GenBank nucleotide accession numbers of virus genomes or genome segments were retrieved for 4,407 virus isolates (corresponding to 3,106 virus species) with invertebrates and/or vertebrates host annotation from the ICTV Virus Metadata Resource VMR MSL38 v2 ^43^. From the viral genomes, we extracted 71,269 protein records encoded by viral protein-coding genes annotation on GenBank. Since many viral genomes encoded polyproteins, we also relied on GenBank annotation to retrieve 4,070 mature peptides from 489 records annotated as polyproteins and 11,786 protein regions from 2,087 proteins with length greater or equal to 2,000 amino acids (aa). 360 proteins had both peptide and region annotation. All records had protein sequence lengths greater than 10 aa. In total, 69,053 proteins without peptide or region annotation, 11,786 protein regions, 4,070 mature peptides, and 253 shortest polyproteins with peptide and/ or region annotation were used for protein structure prediction.

### Protein structure prediction with ColabFold

Protein models for all 85,162 records were predicted using LocalColabFold v.1.5.2 with default settings ^30^. Multiple sequences alignments (MSAs) were constructed for each record using MMseqs2 v.15.6f452 ^44^, and colabfold_envdb_202108, pdb100_230517, uniref30_2302 databases installed locally. Five models were inferred for each record, models were ranked based on mean predicted local-distance difference test (pLDDT) score. The top ranked model was subjected to constrained relaxation by gradient descent in the Amber force field with the following settings: max_iterations 2,000; tolerance 2.39; stiffness 10.0. The relaxed models were used for structural analysis.

### Protein structure prediction with ESMFold

We also predicted 84,964 protein models for 69,043 proteins without peptide or region annotation, 4,070 mature peptides, 11,767 protein regions, and 84 proteins with peptide and/or region annotation using ESMFold module of ESM-2 v.1.0.3 ^24^ with default settings. For records with protein length greater than 1,236 aa following settings were used: --max-tokens-per-batch 1 --chunk-size 128. Due to memory constraints on the GPU, we were unable to predict models for records with protein length greater than 2,840 aa. ESMFold models were subjected to relaxation in the Amber force field as explained above.

### Estimation of structural coverage expansion

To estimate the expansion of structural coverage relative to experimentally determined structures available in the PDB ^21^, and to consider the influence of PDB training data on prediction accuracy, we performed a sequence similarity search of 71,269 protein records against the PDB 2024-02-20 using MMseqs2 v.15.6f452 ^44^ easy-search command with the following sensitivity settings: -s 7.5 --max-seqs 100,000 -e 10^-3^. To calculate the percentage of residues covered by PDB structures, only PDB hits with sequence identity greater or equal to 0.95 were retained. To calculate the percentage of residues covered by homologues from the PDB, PDB entries released after 2018-04-30 or 2020-05-01, training data cut-offs for AlphaFold2 ^23^ and for ESMFold ^24^ respectively, were filtered out, only PDB hits with sequence identity of 30% or greater were retained.

### Clustering procedure

79,036 ColabFold and 6,126 ESMFold models with the highest pLDDT score per protein record were combined into a non-redundant dataset for the downstream structural analysis. ESMFold models were used as representatives of protein records if they had mean pLDDT score greater than 50 and this score was greater than the mean pLDDT score of a corresponding ColabFold structure. First, we clustered 85,162 protein records based on their sequence similarity using MMseqs2 v.15.6f452 ^44^ easy-cluster command with minimum sequence identity of 50% and bi-directional coverage of 90% (--min-seq-id 0.5 --cov-mode 0 -c 0.9 -e 10^-3^ --cluster-mode 0). Second, we defined structural representatives for 33,151 MMseqs2 clusters by choosing the member with the highest model pLDDT score. We clustered 33,151 structure representatives based on structural similarity using Foldseek v.9.427df8a ^45^ easy-cluster command with no minimum sequence identity but bi-directional coverage of 90% (--min-seq-id 0 --cov-mode 0 -c 0.9 -e 10^-5^) producing 19,067 structural clusters.

### Structural and functional homogeneity of clusters

To estimate structural homogeneity within each cluster we extracted LDDT and alignment TM scores from a reciprocal Foldseek v.9.427df8a ^45^ easy-search of the non-redundant dataset with next settings: --exhaustive-search -e 0.1. For each cluster we calculated the median LDDT and TM score of the alignments between the cluster representative and other cluster members. To estimate the consistency of function annotation within clusters we relied on a sequence-based Pfam annotation acquired using InterProScan v.5.69-101.0 ^46^ with default settings. For each protein record we retained a functional annotation with the highest sequence coverage and calculated the percentage of annotated records that share the predominant Pfam annotation for each cluster containing at least two annotated members.

### Structure similarity network analysis

Similarly to MMseqs2 clusters, we defined structural representatives for 19,067 Foldseek clusters by choosing the model with the highest pLDDT score from each cluster. We performed a structural comparison of the representatives using Foldseek v.9.427df8a ^45^ easy-search command with default settings and applied the results to construct a structure similarity network using NetworkX v.3.3 ^47^. Only structural representatives with pLDDT score above 50 and at least one connection with Foldseek E-value below 10^-3^ were retained, producing a graph with 7,812 nodes and 37,751 edges. The graph was visualized using NetworkX spring_layout() function with k of 0.3 and 500 iterations.

### Identification of hallmark proteins and fusion glycoproteins

We started with Foldseek v.9.427df8a ^45^ easy-search against the non-redundant dataset with -e 10^-5^ --max-seqs 10,000 settings using a set of experimentally determined structures as probes: poliovirus RNA dependent RNA polymerase (PDB ID: 4R0E ^48^), human immunodeficiency virus reverse transcriptase (1HMV ^49^), Phi29 DNA polymerase B (2PY5 ^50^), circovirus rolling-circle replication endonuclease (8H56 ^51^), papillomavirus hexameric superfamily 3 helicase (5A9K ^52^), poliovirus VP3 protein (8E8R ^53^) as single jelly roll capsid protein, human adenovirus 5 hexon protein (6B1T ^54^) as double jelly roll capsid protein, varicella zoster virus HK97 capsid protein (6LGL ^55^), respiratory syncytial virus F protein (6APB ^56^) as a class I fusion glycoprotein, spondweni virus E protein (6ZQI ^57^) as a class II fusion glycoprotein, human cytomegalovirus gB protein (7KDP ^58^) as a class III fusion glycoprotein. To test if clustering of structures and structure similarity network analysis increases the number of hits, we performed another Foldseek easy-search using the same list of probes and search parameters but this time against a set of 19,067 structural representatives. We then increased the number of structurally similar clusters by adding clusters that have a network connection (E-value 10^-5^ and 90% coverage of the Foldseek hit) to clusters found using Foldseek. Finally, the list of hits was expanded to all members of the clusters identified with Foldseek and the structural network.

### Expansion of functional annotations

To demonstrate how structural information can be used to expand functional annotation, first we performed a sequence-based annotation of 85,162 proteins using InterProScan v.5.69-101.0 ^46^ with default settings. We focused on annotations from Pfam ^59^, Superfamily ^60^ and Gene3D ^61^ databases because they provided the highest number of annotated proteins: 55,869, 28,431 and 22,336 proteins respectively. From each database we retained only one functional annotation for each protein based on the sequence coverage; shorter annotations were neglected in this analysis. To annotate proteins with unknown function within annotated clusters we used structural clusters: for each cluster we calculated frequencies of annotations coming from annotated members and propagated the predominant annotation to members with unknown function. To annotate clusters with unknown function we used structural network: we calculated frequencies of annotations coming from structural representatives of connected clusters and propagated the predominant annotation to the clusters with no annotation. In case of annotation expansion using the structural network, only connections with Foldseek E-value below 10^-3^ and 90% coverage of InterProScan annotation were considered. Expansion of Gene Ontology (GO) terms ^62^ was done in the same way based on GO IDs associated with Pfam annotations described above.

### Structural comparison to AlphaFold structural database

We compared 19,067 structural representatives of Foldseek clusters to protein models in the AlphaFold Structural Database ^25^ using Foldseek v.9.427df8a ^45^ easy-search option with E-value of 10^-3^. For each structural representative we retained the hit with the lowest E-value.

### Comparison to alternative repositories of viral protein structure predictions

We systematically compared Viro3D to the dataset generated by Nomburg et al. and to the Big Fantastic Virus Database (BFVD) ^28,29^. We first used Foldseek searches to assess overlap with Viro3D. Here, protein models were considered shared if they had a Viro3D Foldseek hit with an E-value lower than 1e^-5^, sequence identity equal or greater than 95% and query coverage equal or greater than 95%. Next, we examined structure predictions from a panel of priority human viral pathogens. Using taxonomic identifiers to find the relevant models, we compiled structure sets for each pathogen from Viro3D, Nomburg et al. and BFVD. Using protein sequence to identify matching models (95% sequence similarity), we evaluated coverage of proteins (relative to Viro3D) and the confidence of structure prediction (assessed using pLDDT values).

### Building a structural-similarity map displaying all viral species

We first performed all-vs-all Foldseek comparison of the entire non-redundant structure set (85,162 models), therefore surveying global structural-similarity. Using this, for any given virus’ proteome (i.e. collection of structures), we extracted the lowest E-value against each of the 85,162 models. If no E-value was present for a given virus-model pair, we substituted an arbitrary high value of 10. This resulted in a matrix where each of the 4,407 viruses is represented by 85,162 structural similarity scores. We then used principal component dimensionality reduction to group viruses based on the structural-similarity of their respective proteomes.

### Phylogenetic analysis of the Class I fusion glycoprotein

The complete set of Class I structures, identified with Foldseek and the structural network, were aligned using two approaches: i) FoldMason v.99fbda50e6c296c9fdf15d05fefedbb91f4efe84 ^63^ (easy-msa –report-mode 1) and ii) the famsa3di method described in Puente-Lelievre et al. (2024) ^64,65^, producing corresponding 3Di and amino acid alignments. The alignments were also trimmed using trimAl with a gap threshold of 35% ^66^, consistent with Mifsud et al. (2024) ^67^. Subsequently, the phylogenetic relationships were determined using IQ-TREE v.2.3.6 ^68^ using the four 3Di alignments (FoldMason, FoldMason trimmed, famsa3di, famsa3di trimmed). Suitable substitution models were tested by BIC using ModelFinder ^69^ and the custom 3DI substitution matrix ^65^ with empirically counted frequencies from the alignment and accounting for rate heterogeneity with a six category FreeRate model (3DI+F+R6) was the best (or near best) model for all 3Di alignments and used consistently for all tree inferences. To inform phylogenetic reconstruction with amino acid homology we also performed partitioned model inferences where one partition corresponds to the 3Di alignment, using the 3DI+F+R6 substitution model and the other to the amino acid alignment, using a LG+F+R6 substitution model. Branch support was assessed using 10,000 replicates of the ultrafast bootstrap approximation method ^70^. Phylogenetic trees were visualised and prepared for publication with the ggtree R package v3.10.1 ^71^. All accompanying structure models were visualised and prepared for publication with UCSF ChimeraX ^72^.

### Development of the Viro3D web-resource

The frontend of Viro3D was built using React and Typescript providing a dynamic user interface. To enhance specific features, we integrated PDBE-Molstar Viewer to provide an interactive 3D rendering of protein structures (https://github.com/molstar/pdbe-molstar); Sodaviz (https://github.com/sodaviz) to allow for browsing protein structures across the virus genome; KonvaJS to construct the interactive map of viruses based on structure similarities (https://github.com/konvajs/konva). The Viro3D backend was developed using FastAPI to allow for programmatic access to the data. Biopython was integrated for leveraging the NCBI BLASTp command-line wrapper to enable searching by protein sequence. The MongoDB Community Edition was used to store the data.

## Acknowledgments

This work was funded by UK Medical Research Council (MRC) support to the MRC-University of Glasgow Centre for Virus Research (CVR Integrative Viral Genomics and Bioinformatics Platform: MC_UU_00034/5, CVR Preparedness Platform: MC_UU_00034/6 and CVR Structure-to-Function of Virions Programme: MC_UU_00034/1). This research is jointly funded by the MRC and the Foreign Commonwealth and Development Office (FCDO) under the MRC/FCDO Concordat agreement. J.G. is also supported by a Wellcome Trust and Royal Society Sir Henry Dale Fellowship (107653/Z/15/Z) and an Emerging Leaders Prize from the Medical Research Foundation. U.L. is supported by a PhD studentship from the Darwin Trust of Edinburgh. We thank Prof. Emma Thomson and Prof. Massimo Palmarini for their support of the Viro3D project.

---

**Figure S1.**
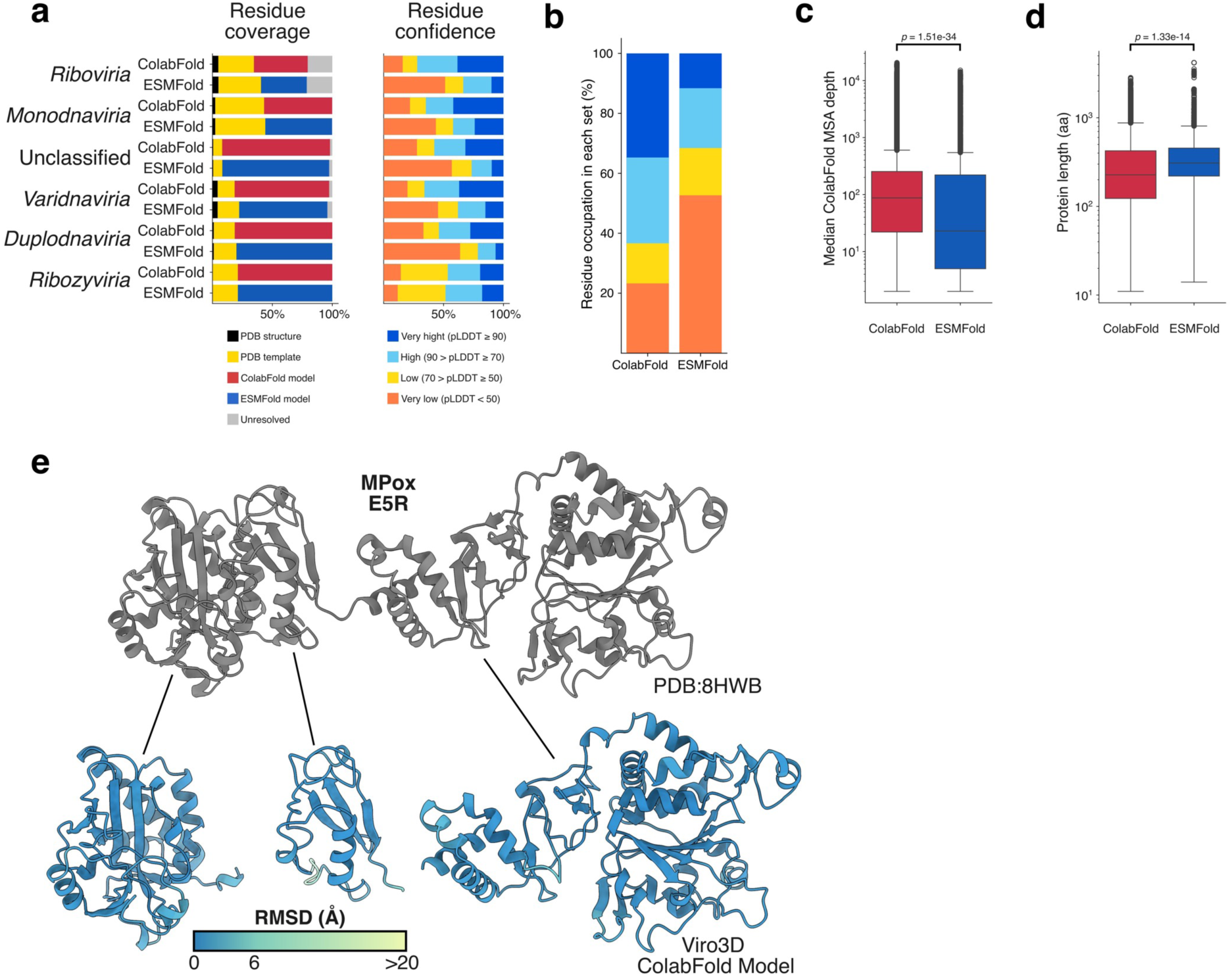
Expanding structural coverage of the human and animal virosphere. **a.** On the left, percentage of residues covered by ColabFold (red) and ESMFold (blue) models in contrast to percentage of residues covered by PDB structures (sequence identity >= 95%, black) and PDB templates (sequence identity >= 30%, yellow) for each viral realm. On the right, confidence of residues modelled by ColabFold and ESMFold based on pLDDT score per viral realm. **b.** Percentage of ColabFold and ESMFold residues with very high (dark blue), high (light blue), low (yellow) and very low (orange) pLDDT score. **c.** Distributions of median ColabFold MSA depth for records where ColabFold model has a higher pLDDT score (red box) and ESMFold models has a higher pLDDT score (blue box). The difference between MSA depth is significant based on the t-test results (t-statistic: -12.26; *p*-value: 1.51e-34). Box represents the interquartile range, the line inside the box represents the median value. Whiskers extend from the box to the smallest and largest values within 1.5 times the interquartile range. Outliers and are plotted as individual points. **d.** Distributions of protein length for records where ColabFold model has a higher pLDDT score (red box) and ESMFold models has a higher pLDDT score (blue box). The difference between protein length is significant based on the t-test results (t-statistic: -7.70; *p*-value: 1.33e-14). **e.** Mpox E5R protein, as shown in Figure 1g, but with the three constituent individual domains color-coded by RMSD (Å) after their respective rigid-body alignment with the experimental structure (grey). This indicates that whilst their relative positions may diverge from the experimental structure (Figure 1g), the individual domains are accurately predicted.

**Figure S2.**
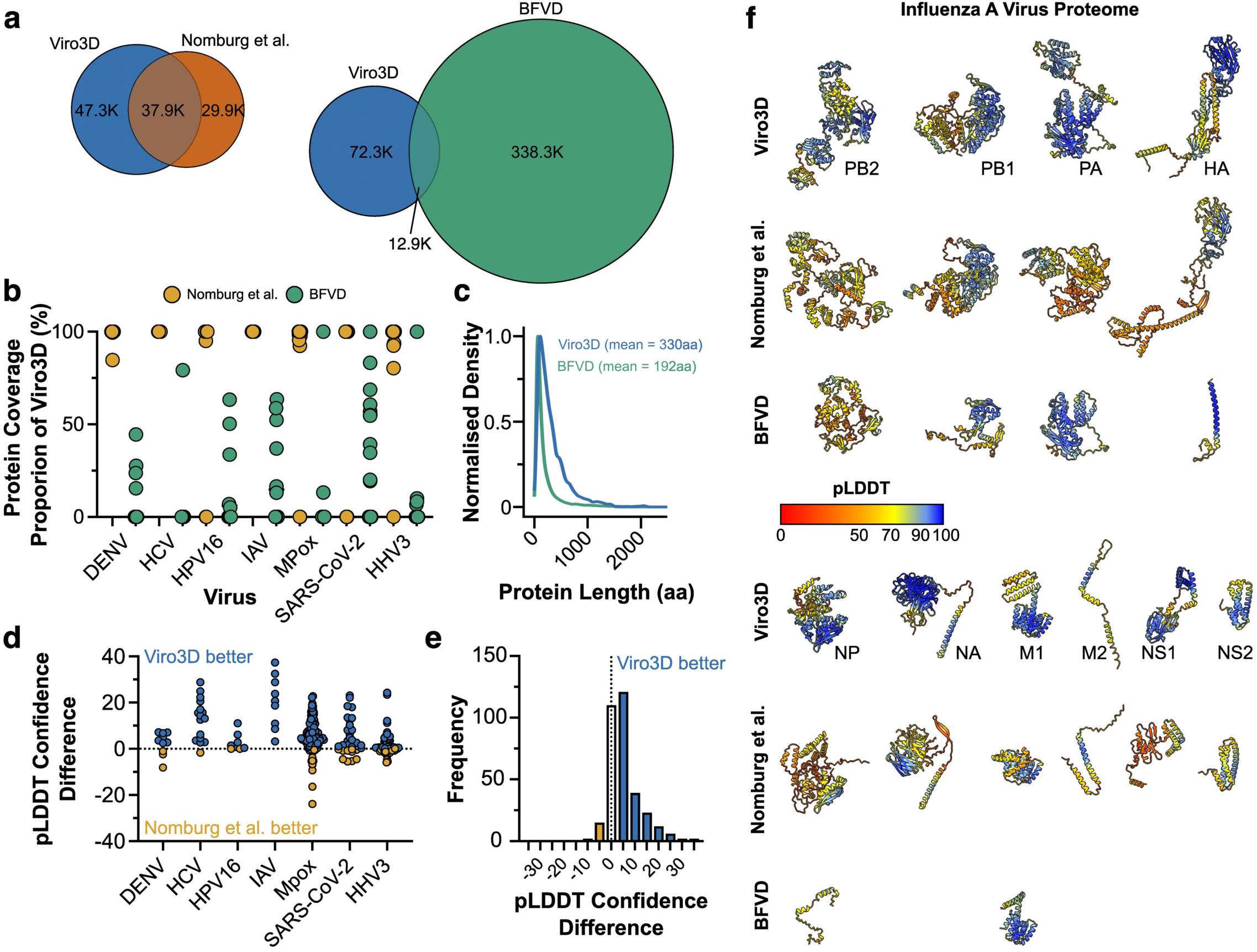
Comparison of Viro3D with alternative repositories of viral protein structure predictions. Viro3D was systematically compared to the Nomburg et al. dataset and the Big Fantastic Virus Database. **a.** Foldseek was used to find matching protein structures with the following threshold parameters: E-value lower than 1e^-5^, sequence identity ≥95% and query coverage ≥95%. Venn diagrams demonstrate overlap in datasets. We also compared proteome level structure sets from each database for example human pathogens: dengue virus (DENV), hepatitis C virus genotype 1a (HCV), human papilloma virus 16 (HPV16), influenza A virus PR8 (IAV), Mpox, severe acute respiratory syndrome coronavirus-2 (SARS-CoV-2) and human alphaherpes virus 3 (HHV3). **b.** Each dot displays the coverage of individual proteins in the alternative datasets (as a proportion of the Viro3D model). **c.** Distribution of protein lengths (in amino acid residues) for all entries in Viro3D and BFVD. **d.** Proteome level comparisons, as in b., of structure prediction confidence (pLDDT) in matching models from Viro3D and Nomburg et al.. Data is expressed as difference in pLDDT scores, with positive values indicating a higher score in the Viro3D model. **e.** Histogram of pLDDT difference for the cumulative set of structures shown in d. (n= 334 models). **f.** Predicted Viro3D structures for IAV proteome alongside the best matching counterparts from Nomburg et al. and BFVD. Ribbon diagrams are colour-coded by pLDDT confidence as denoted in the key.

**Figure S3.**
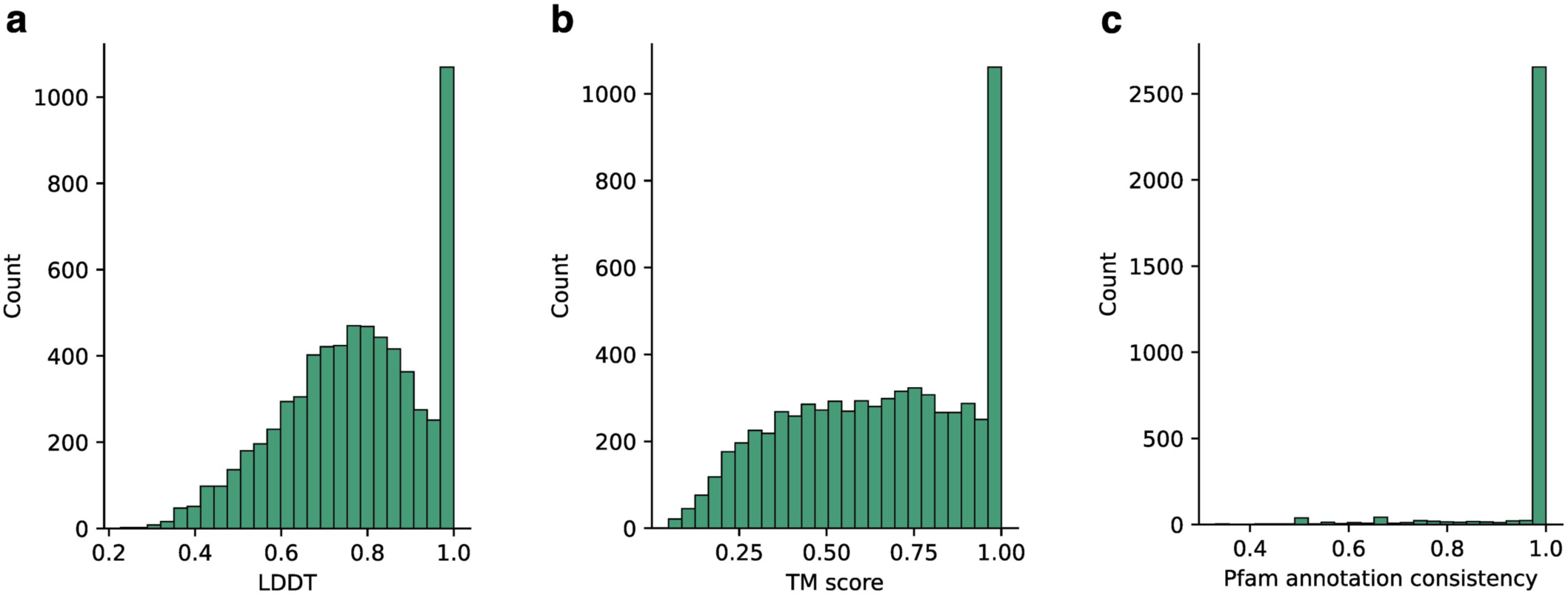
Homogeneity of structural clusters. **a.** Distribution of average LDDT scores across 19,067 structural clusters. LDDT provides a measure of local structural similarity, with identical structures giving a values of 1, the median LDDT score is 0.78. **b.** Distribution of average TM scores across structural clusters. TM score give a measure of global structural similarity, again identical structures give a value of 1, the median TM score is 0.66. **c.** Distribution of Pfam function annotation consistency across structural clusters. We would expect proteins clustered by structure to have the same function, 95% of annotated clusters have Pfam consistency of 100%.

**Figure S4.**
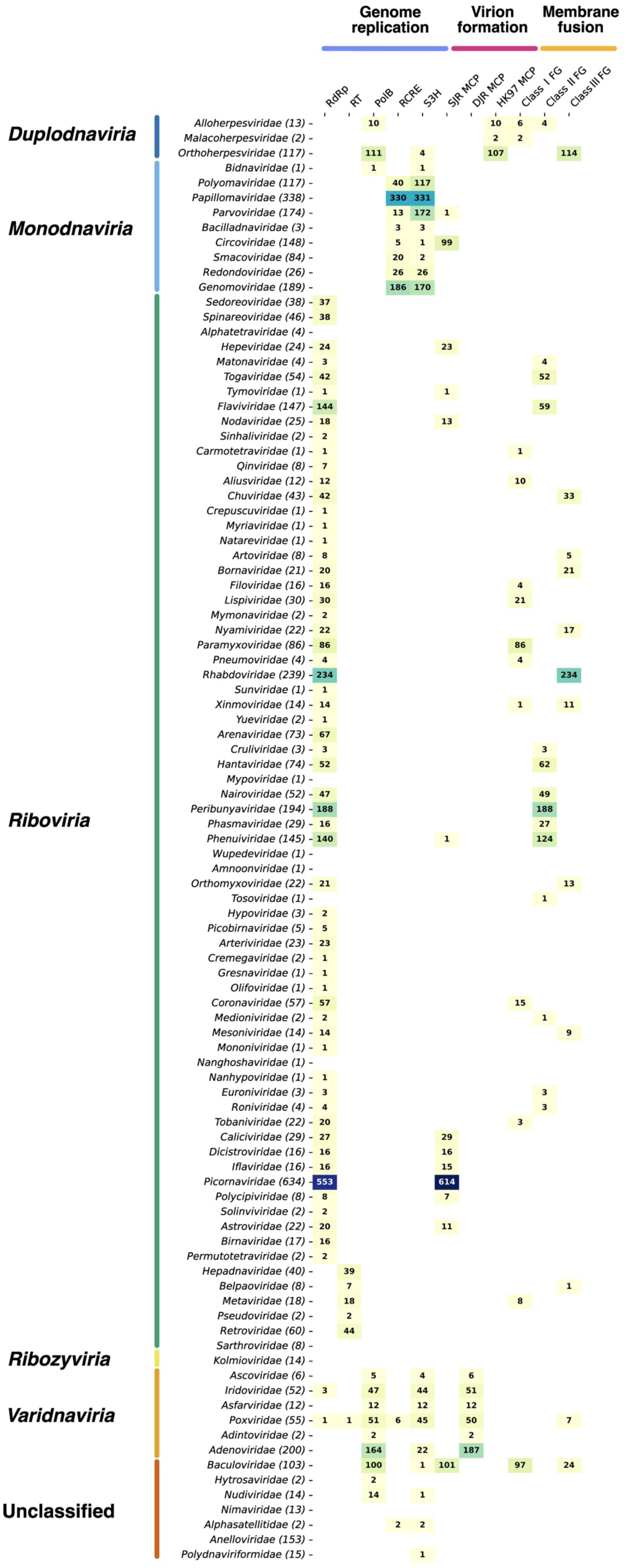
Mapping the distribution of hallmark viral proteins across the human and animal virosphere. Number of viruses with structural homologues to hallmark viral proteins, organised by virus family. Structural homologues were found using a Foldseek search against the structure similarity network (Figure 2) using individual experimental protein structures as references: RNA dependent RNA polymerase (PDB ID: 4R0E), Reverse transcriptase (1HMV), DNA polymerase B (2PY5), Rolling-circle replication endonuclease (8H56), Superfamily 3 helicase (5A9K), Single jelly roll capsid protein (8E8R), Double jelly roll capsid protein (6B1T), HK97 capsid protein (6LGL), Class I fusion glycoprotein (6APB), Class II fusion glycoprotein (6ZQI), Class III fusion glycoprotein (7KDP).

**Figure S5.**
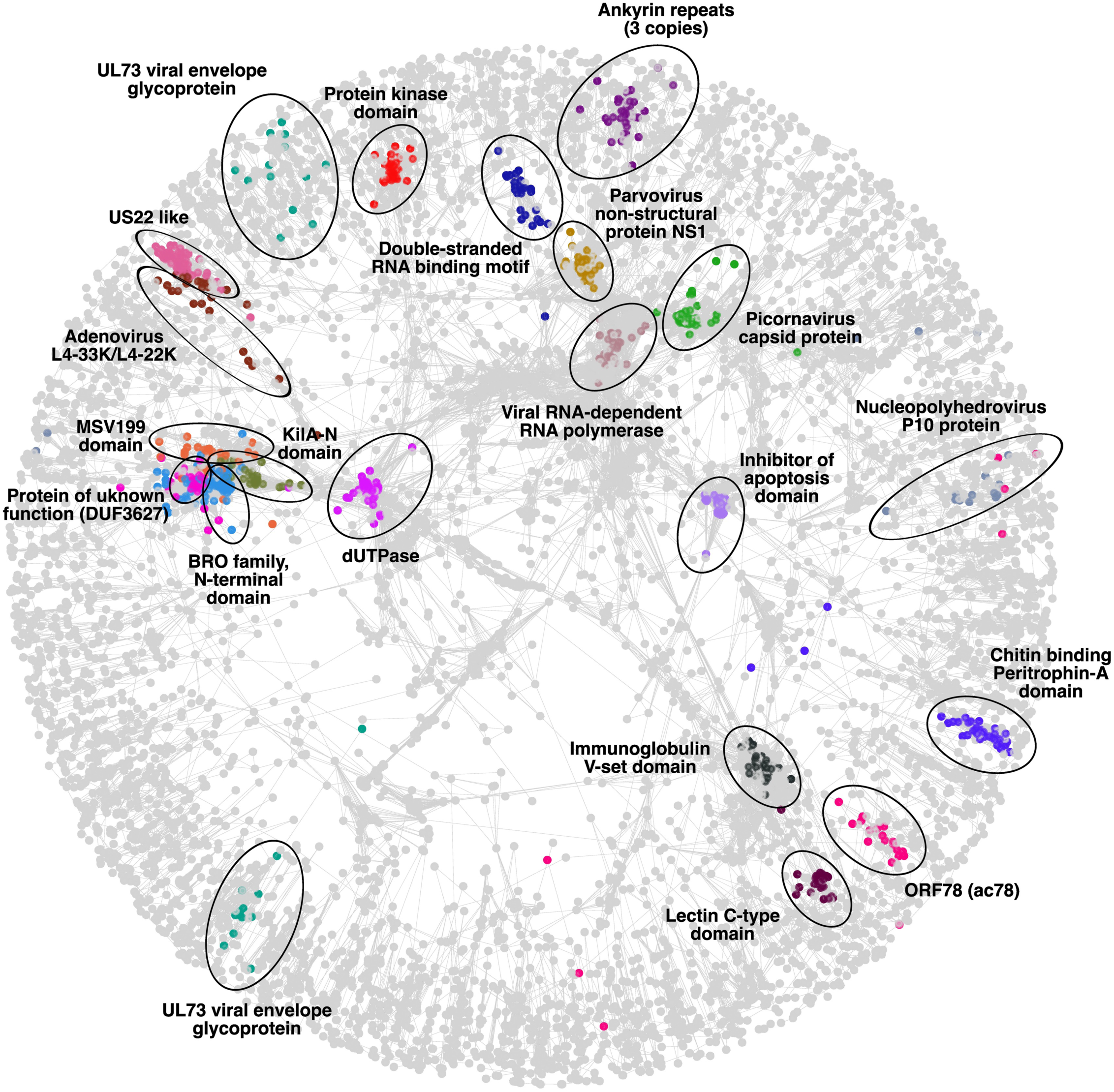
Functionally annotated structure similarity network of viral proteins. Each node represents a cluster. Edges connect clusters that share structural similarity. Clusters that possess the twenty most frequent Pfam annotations are highlighted in different colours.

**Figure S6.**
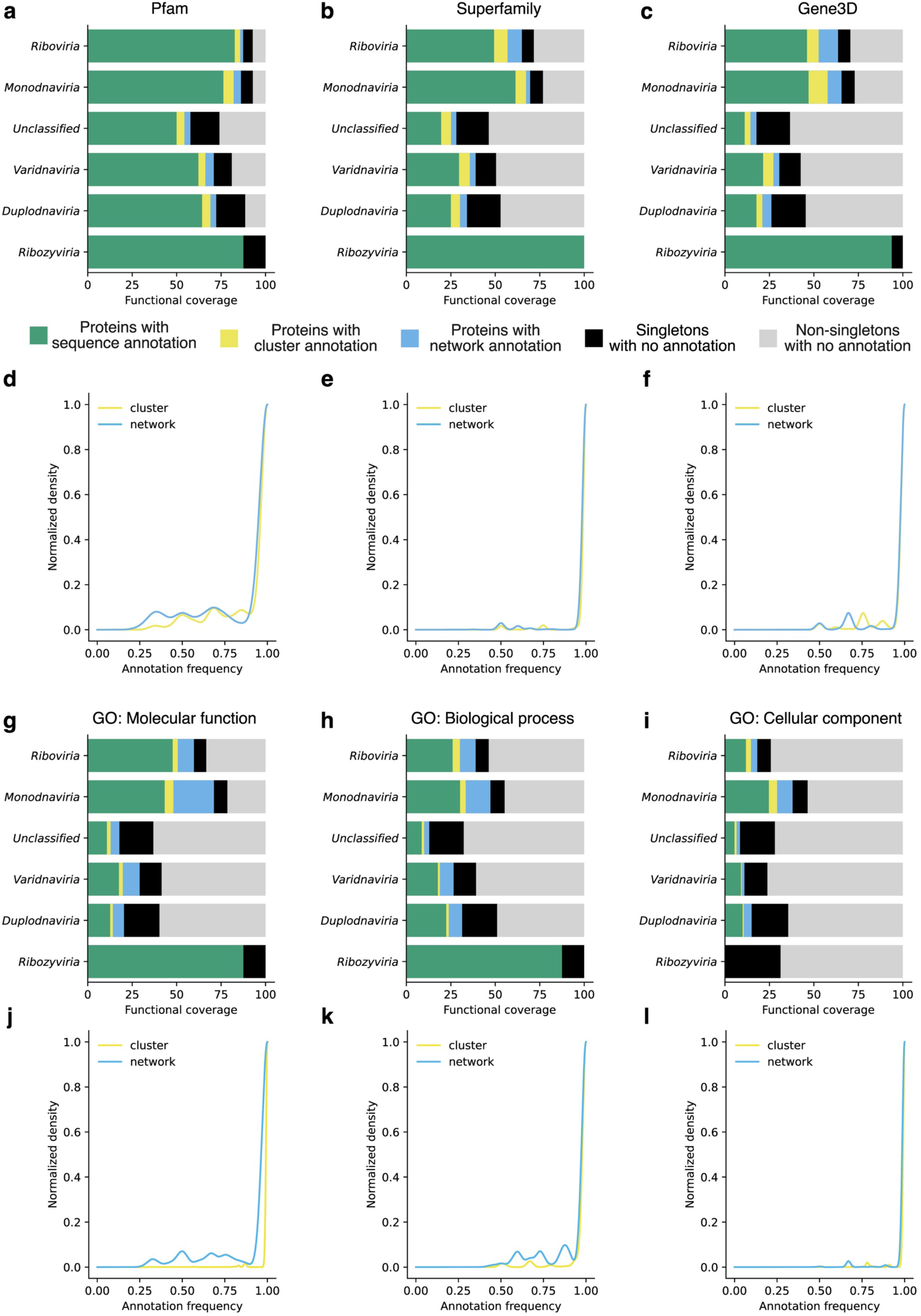
Propagated protein annotations by viral realm. **a-c.** and **g-i.** Percentage of protein records in each viral realm that possess functional annotation based on InterProScan (green), structural cluster expansion (yellow), structural network expansion (light blue). Percentage of protein records that do not have a functional annotation are black (if they belong to singleton clusters) or light grey (if they belong to non-singleton cluster). **d-f.** and **j-l.** Distribution of the frequencies of propagated annotations. Functional annotations propagated using structural clusters are in yellow, annotations propagated using structural network are in light blue.

**Figure S7.**
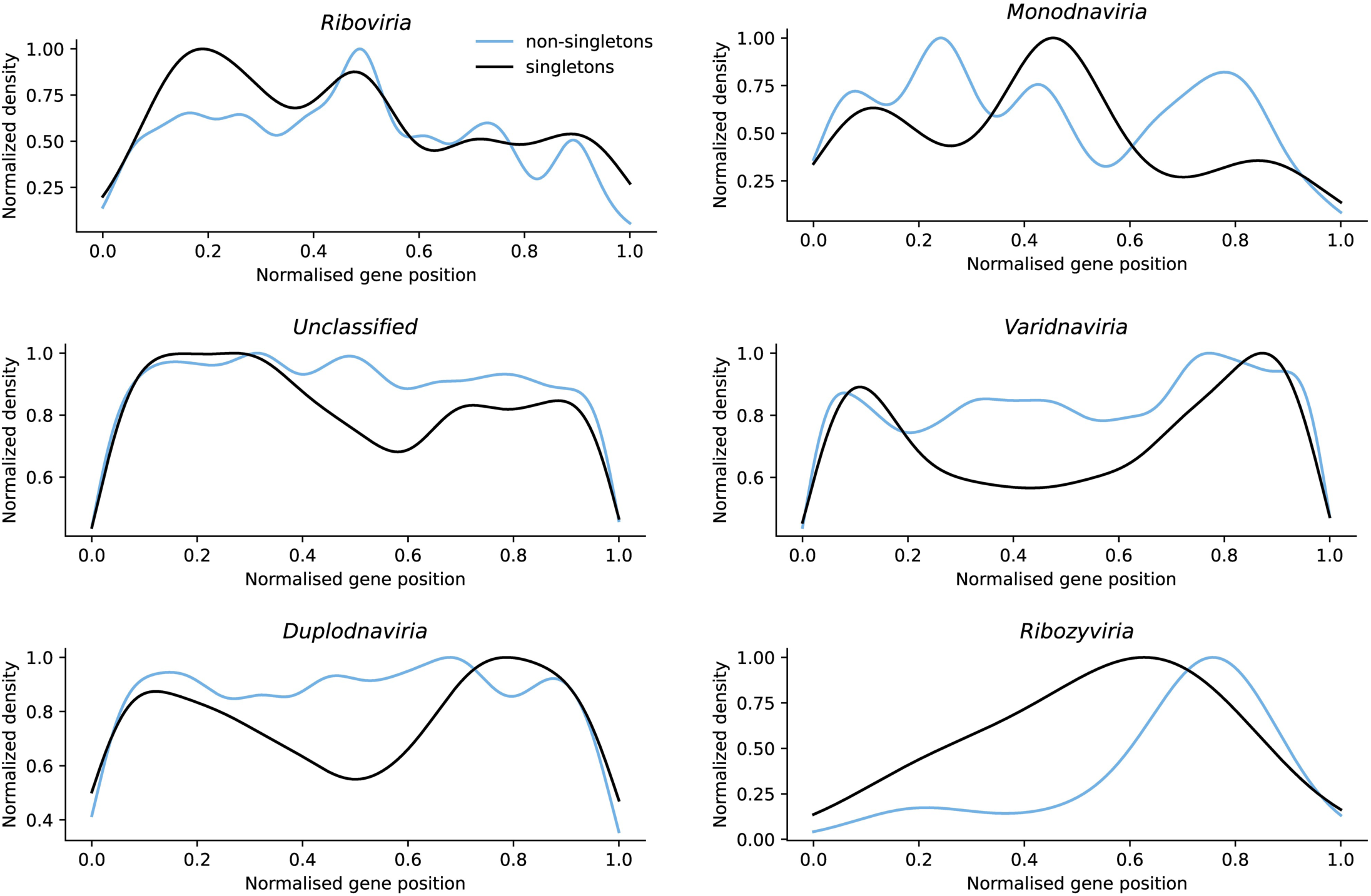
Genomic positions of singleton (unique) and non-singleton protein-coding genes by viral realm. Proteins that belong to singleton clusters are in black, protein that belong to non-singleton clusters are in light blue.

**Figure S8.**
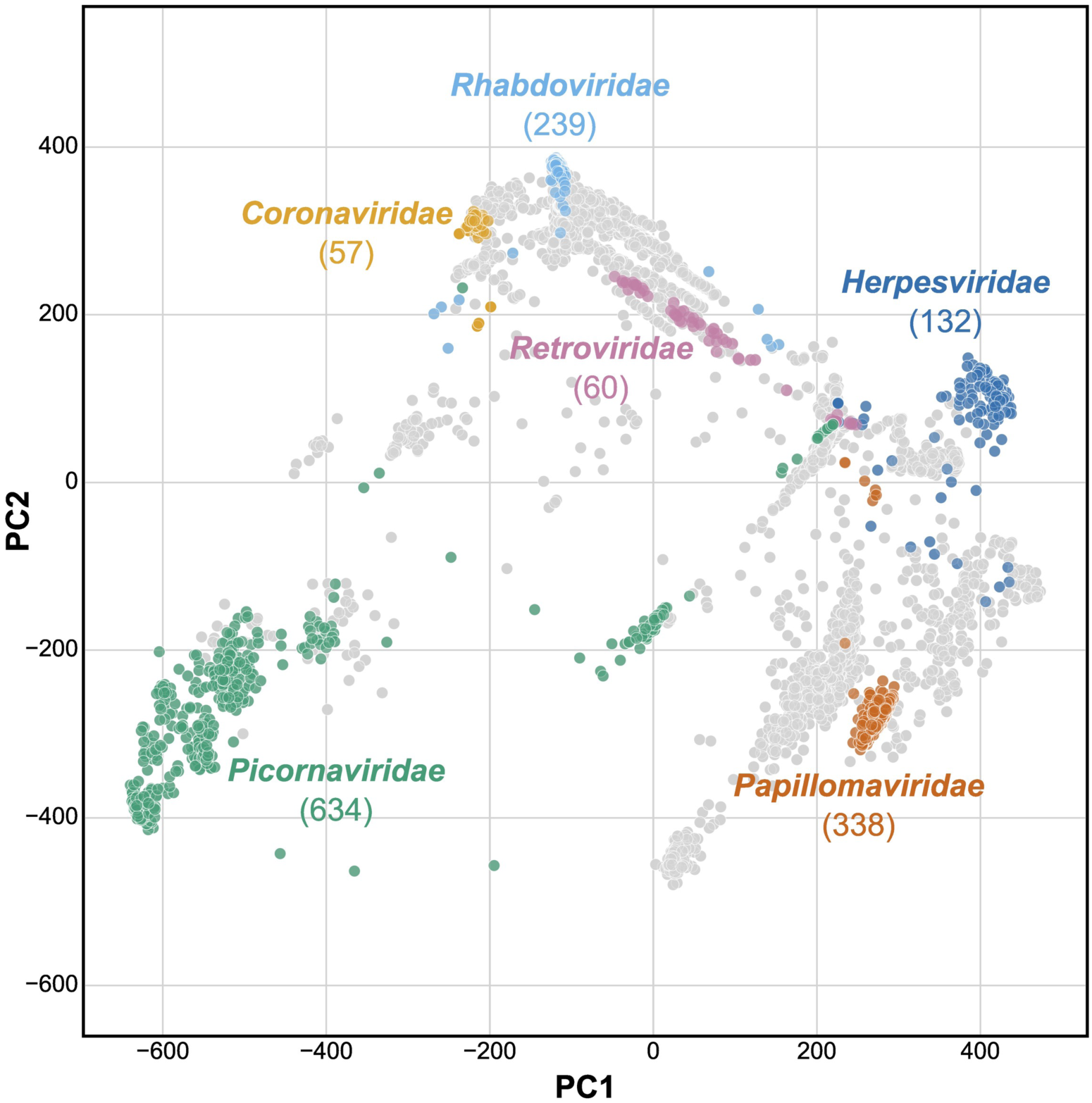
Grouping viruses by structural similarity recapitulates taxonomy. Structure-informed map of the human and animal virosphere, each data point represents a virus in the Viro3D database (see Methods). Coloured data points show example viral families, which mainly produce coherent clusters within the structure-similarity map. Values in parentheses indicate the number of viruses within each family.

